# Adaptation of a microfluidic qPCR system for enzyme kinetics studies

**DOI:** 10.1101/2020.09.18.303248

**Authors:** Elzbieta Rembeza, Martin KM Engqvist

## Abstract

Microfluidic platforms offer a drastic increase in throughput while minimizing sample usage and hands-on time, which makes them important tools for large-scale biological studies. A range of such systems have been developed for enzyme activity studies, although their complexity largely hinders their application by a wider scientific community. Here we present adaptation of an easy-to-use commercial microfluidic qPCR system for performing enzyme kinetics studies. We demonstrate functionality of the Fluidigm Biomark HD system (the Fluidigm system) by determining kinetic properties of three oxidases in a resorufin-based fluorescent assay. The results obtained in the microfluidic system proved reproducible and comparable to the ones obtained in a standard microplate-based assay. With a wide range of easy-to-use, off-the-shelf components, the microfluidic system presents itself as a simple and customizable platform for high-throughput enzyme activity studies.

**Graphical abstract:** 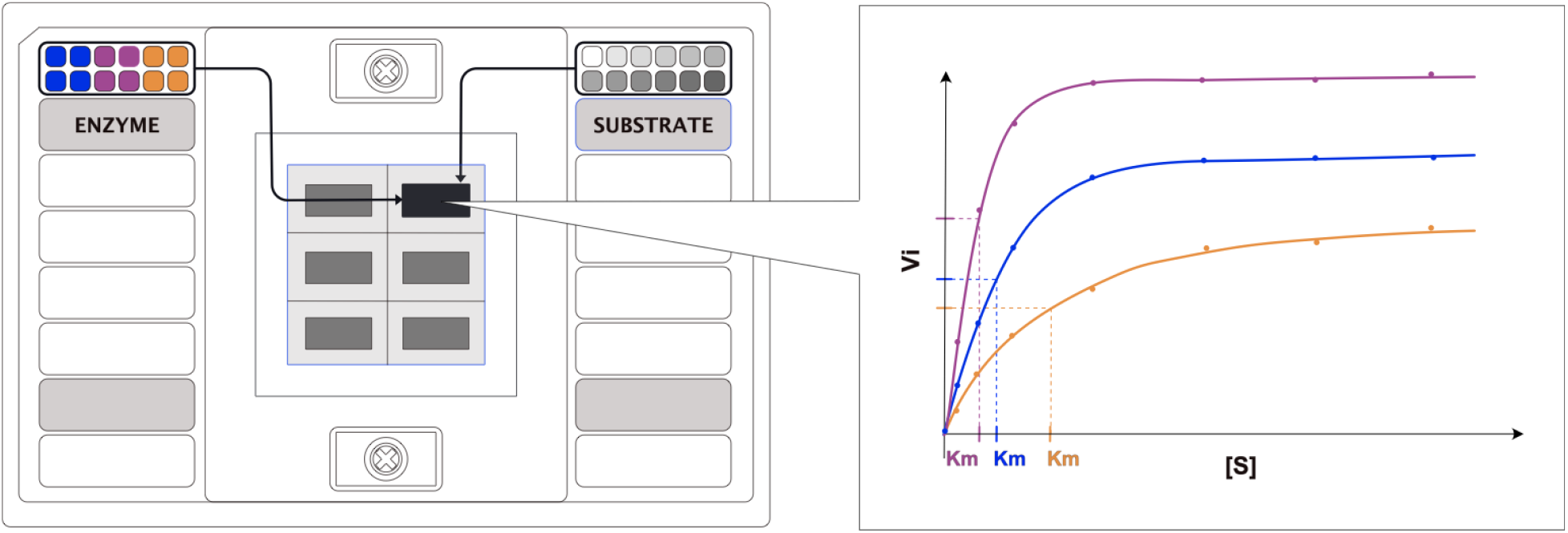

## Introduction

In the past decade we have been witnessing a drastic increase of genomic data deposited in public databases, which provides the potential for discovery of new enzymatic activities and deeper understanding of the known ones ^1–3^. However, current experimental methods cannot keep up with characterisation of novel genes, resulting in a chronic lack of validated data in public databases ^4^ Currently, the most common methods for in-depth enzyme characterisation rely on manual preparation of enzyme-substrate mixtures in a microplate, which is time-consuming and requires large amounts of protein and substrate. This usually results in a limited range of conditions and substrates being tested and hinders the exploration of enzymatic potential of a protein. Number of high-throughput methods have been developed for both end-point and kinetic measurements of enzyme activity ^5–7^, however, their adaptation by a wider scientific community proves difficult, as it requires specialized knowledge and access to microfluidic manufacturing facilities ^8^. A simple method for performing enzyme kinetic studies in readily available high-throughput off-the-shelf devices would help to address this problem.

Although no high-throughput microfluidic devices for measuring enzyme kinetics are commercially available, such instruments were developed for real-time qPCR applications ^9,10^. One such system was established by Fluidigm Corporation, where gene expression analysis is carried in integrated fluidic circuit chips, with a setup that offers an increase in throughput and reduction in sample volumes, yet remains user-friendly ^11^. In the Fluidigm system samples and reagents are first pressure-loaded into nanoliter-sized reaction chambers of the chip, which is then transferred to a real-time PCR instrument designed to thermal cycle the microfluidic chips and image the data in real time. The Fluidigm ecosystem, including a number of different chips and a range of available fluorescent filters (Supplementary table 1), indicates its potential for novel applications, in addition to qPCR. In this work we show that a qPCR microfluidic system can be easily adapted for enzyme kinetic studies with improved throughput and decreased sample usage over conventional methods.

## Results

### Characterisation of experimental platform: microfluidics qPCR device for enzymatic assays

To assess the suitability of Fluidigm system for enzyme activity screening, we chose a FlexSix Gene Expression IFC chip which contains six partitions, each with 12 wells on the assay side and 12 wells on the sample side each (Figure 1A). The chip provides a medium range of throughput: from 144 up to 864 reactions per chip, depending on how samples are routed in the chip. As a model enzyme we selected the hydrogen peroxide producing enzyme lactate oxidase (LOX, EC 1.1.3.2). Activity of lactate oxidase can be detected by a simple coupled fluorescent assay, in which a non-fluorescent probe reacts with hydrogen peroxide to produce a fluorescent product, resorufin (Supplementary figure 1).

**Figure 1.**
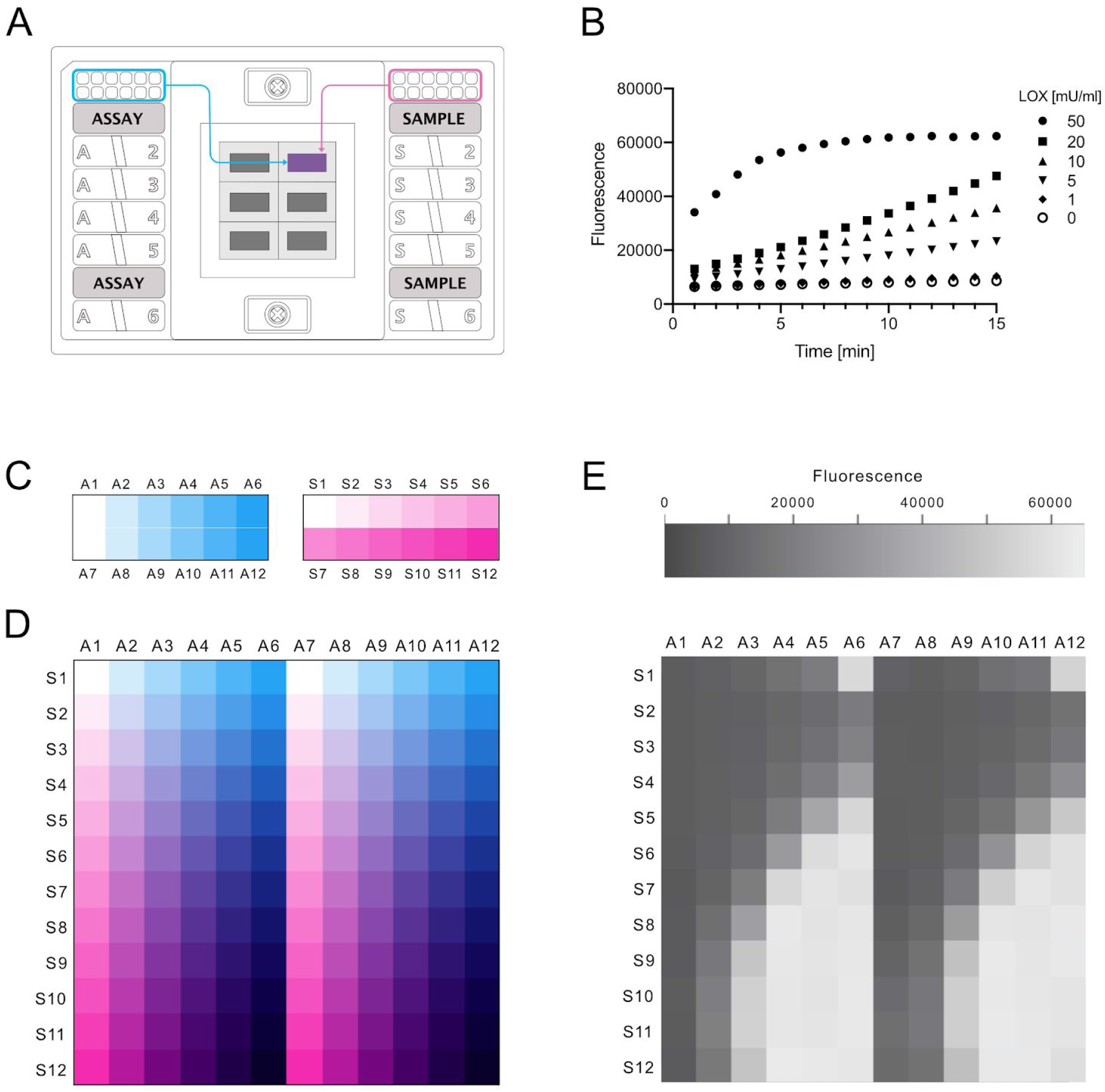
Exploration of the Fluidigm gene expression system as a platform for enzyme activity measurements. (A) Schematic representation of a FlexSix microfluidic chip. (B) Example of obtained initial reaction rates of lactate oxidase in reaction with lactate in buffer composition with 0.01% Triton-X-100 and Tween 20 and addition of BSA. (C) Experimental setup for measurements of enzyme kinetic parameters in the Biomark HD system. Schematic overview of enzyme loading with 6 different concentrations (blue) in the “assay” input wells and loading of substrate with 12 different concentrations (magenta) in the “sample” input wells. (D) Schematic overview of the sample/assay gradient that results from the all vs all mixing that occurs in the FlexSix chip. All enzyme concentrations are mixed with all substrate concentrations. In the illustrated scheme two technical duplicates of each enzyme concentration are run. (E) Example of resorufin fluorescence intensity values recorded in the run. Here for lactate oxidase at cycle 60.

We first tested over what range resorufin fluorescence was linear in the Fluidigm system by measuring a range of resorufin concentrations. Our results indicated that it is possible to capture linear response for up to 5 μM of resorufin, which is comparable to data obtained in a microplate reader (Supplementary figure 2). Next, we asked whether enzyme activity could be captured in the system, and measured the increase of fluorescence over time for different enzyme concentrations with 5 mM lactate. To lower adhesion of protein to the chip’s channels and minimize protein precipitation, we tested two concentrations of nonionic detergents in a buffer, in presence or absence of BSA. Additionally, we used fluorescein dye as a loading control to inspect whether the mixing of the sample is consistent throughout the chip. The results showed that not only were we able to detect the activity of the enzyme, but also to capture its initial reaction rates (Figure 1B). Addition of BSA did not influence the results, while higher concentration of detergents provided a more consistent signal readouts throughout replicates but resulted in much higher overall background fluorescence (Supplementary figure 3). Readouts of the fluorescent signal from the fluorescein loading control indicate equal sample loading throughout the chip, with higher detergent concentrations leading to more reproducible loading (Supplementary figure 4). Overall, the results of the initial screen show the potential of the Fluidigm system for measuring enzyme activity.

### Measurement of enzyme kinetics in the Fluidigm microfluidic system

Next, we asked whether the Fluidigm system is reliable for studying enzymatic Michaelis-Menten kinetics. To test this, we measured initial reaction rates of three hydrogen peroxide producing enzymes: lactate oxidase, glucose oxidase (EC 1.1.3.4) and glutamate oxidase (EC 1.4.3.11). For each of the three oxidases, five different enzyme concentrations were placed in duplicates on the assay side of the FlexSix chip, and 11 substrate concentrations were placed on the sample side (Figure 1C). During the sample loading, in each partition, all enzyme samples were separately mixed with all substrate samples, creating an enzyme-substrate gradient (Figure 1DE). The use of different enzyme concentrations enabled capturing kinetic information in one run only, without the need of previous knowledge of the enzyme activity. Final enzyme concentrations were between 9.6 ng/ml and 6 μg/ml, and final substrate concentrations were between 1.7 μM and 100 mM. To enable rapid analysis of this data we developed R-scripts which automatically identifies the enzyme concentration for which initial rates are linear, fits a linear regression model to those data points, and extracts the slope.

When establishing a new method it is of key importance to validate it. We therefore set out to test the systems reproducibility as well as accuracy. In order to test reproducibility of the system, we repeated the assay for each enzyme three times using three different FlexSix chips. The obtained kinetic parameters of the three replicates were highly similar (Table 1, Supplementary figure 5), indicating outstanding reproducibility of the method. As a validation of the system’s accuracy, we compared the results obtained in the Fluidigm system with the ones obtained from assays performed in a standard 384-well plate with signal recorded in a fluorescent plate reader (Table 1, Supplementary figure 5). The largest difference in the kinetic values between the two methods was observed for the glucose oxidase (microfluidic chip: *K*_M_ = 26 mM, *V*_max_ = 31.67 μmol mg^−1^ min^−1^, microplate: *K*_M_ = 51.3 mM, *V*_max_ = 9.7 μmol mg^−1^ min^−1^), and was a result of a large deviation obtained in the plate reader measurements, most likely due to a manual pipetting error. Overall, the results show that the performance of the Fluidigm system in measuring enzyme kinetics is on par, in terms of data quality, with that of a standard plate system used routinely today.

**Table 1.** Comparison of kinetic values of three oxidases obtained in the Fluidigm system and in a microplate reader. Values represent mean average (+/- standard error of mean; n = 3).

| Enzyme | System | $K_M$ [mM] | $V_{max}$ [$\mu\text{mol mg}^{-1} \text{min}^{-1}$] |
| --- | --- | --- | --- |
| Lactate oxidase | microplate | $1.15 \pm 0.15$ | $34.67 \pm 1.33$ |
| | chip | $1.40 \pm 0.06$ | $25.67 \pm 3.18$ |
| Glutamate oxidase | microplate | $0.20 \pm 0.03$ | $6.93 \pm 0.43$ |
| | chip | $0.24 \pm 0.04$ | $6.63 \pm 0.94$ |
| Glucose oxidase | microplate | $51.30 \pm 17.3$ | $9.70 \pm 1.67$ |
| | chip | $26.00 \pm 4.34$ | $31.67 \pm 3.33$ |

## Discussion

In this work we show that a microfluidic qPCR system can be applied in biochemistry without modification and is suitable for determining enzyme kinetic parameters. Although we have only tested one microfluidic system in this work, the method can likely be adapted to systems from other manufacturers. The Fluidigm system offers the well-known advantages of microfluidic devices over conventional methods: drastic reduction in both reagent volumes used, as well as time and manual handling involved in performing experiments ^5^. Like many microfluidic devices, the Fluidigm system used here comes with the drawback of only being suitable for fluorescent-based assays, which limits the type of enzyme activities that could potentially be monitored. This, however, is becoming less of an issue with an increase of new fluorescent enzyme screens being developed ^12^. The majority of microfluidic approaches for measuring enzyme kinetics rely on creating concentration gradients of enzyme and substrate inside a device ^7^, which is not the case with the system tested here, where some degree of hands-on work is still required in the preparation of sample dilutions. However, the system presented in this work provides a clear advantage over the previously described methods: low system complexity, which allows users to operate it without fluid handling expertise or infrastructure required for microfluidic device fabrication. This is an important advantage, as the lack of easy to use devices is a frequent cause for low adaptation of microfluidics by non-specialists like biochemists ^8^.

A further advantage of the system is that it allows parallelization of many enzyme kinetic measurements at the same time, with the possibility of testing different enzymes, substrates and conditions in a single chip. Availability of many easy to use, off-the-shelf chips makes the Fluidigm system a good platform for different experiment designs for enzyme screening. The system allows both for smaller scale experiments in FlexSix IFCs, as well as large scale, all-versus-all studies using the larger 96.96 IFC; where up to 9216 separate reactions can be performed simultaneously. In our study we present the successful use of one type of chip and two fluorescent probes: resorufin and fluorescein. In order to explore the full potential of the system, additional experiments with a wider range of chips, enzymatic activities and substrates would be required. We believe that adapting a microfluidic gene expression system for measuring enzyme activity can facilitate the growing need for in-depth, yet high-throughput characterisation of enzymes.

## Materials

Flex Six Gene Expression IFC (product number 100-6308) and Control Line Fluid Kit (product number 89000021) were purchased from Fluidigm Corporation, US. All enzymes and chemicals were purchased from Sigma Aldrich, unless stated otherwise.

## Methods

### Fluidigm system setup

The Flex Six Gene Expression IFC was used for all microfluidic experiments. The Flex Six chip has a total of six partitions which can either be run independently or simultaneously. Each partition has a 12×12 format, in which solutions from 12 assay inlets are mixed with solutions from 12 sample inlets in 1:9 ratio. Final reaction volume is 8.9 nl. Samples placed in the assay inlets were prepared as 10x concentrations and solutions placed in the sample inlets were prepared as 1.1x concentrations. Before the first use of each IFC, the chips were primed with Control Line Fluid Kit in Juno system (Fluidigm, US) according to manufacturer’s guidelines. To load the IFC, barrier plugs were removed from the partitions in use, and 3 μl of each assay and each sample were pipetted into their respective inlets. Caution was taken not to introduce air bubbles while pipetting. Next, the IFC was placed in the Juno machine and the “Load Mix Flex Six GE” was run. Directly after the loading script was finished, the IFC was transferred to a Biomark HD machine (Fluidigm, US) and data collection protocol was set up using Fluidigm’s Data Collection Software. After selecting partitions for the run, “Gene Expression” application type was chosen, ROX was picked as “passive reference” (excitation filter: 575 nm, emmision filter: 630 nm) and FAM-MGB as “probe” (excitation filter: 475 nm, emission filter: 525 nm). The ROX channel was used to collect signal from resorufin and FAM-MGB to collect signal from fluorescein loading control, which is the inverse of what is done in a typical qPCR experiment. The chip run protocol was loaded and exposure was set to 0.03 s for ROX and 2 s for FAM. Chip run protocol was created using Fluidigm’s Real-Time PCR Analysis Software: measurements were taken every 30 seconds for 30 minutes, at 25 °C. After the run was finished, the IFC was placed in Juno and “Post Run Flex Six GE” script was run in order to relax the valves.

### Enzyme assays in the Fluidigm platform

All enzyme assays were performed in a buffer containing a final concentration of 20 mM HEPES pH 7.4, 50 μM AmplifluRed, 0.1 U/ml horseradish peroxidase (HRP), 0.01% Tween 20, 0.01% Triton X 100, 0.01 mg/ml BSA, 0.1 μM fluorescein, unless stated otherwise. However, due to how the microfluidic chips function, solutions from assay and sample sides are mixed at a ratio of 1 to 9. One must therefore make use of suitably higher initial concentrations in the assay and sample wells to ensure correct final concentrations inside the chip. 10x concentrated Ampliflu Red, HRP, Tween20, Triton X 100, BSA, fluorescein and enzyme were placed in the assay inlet whereas 1.1x concentrated fluorescein together with substrates were placed in the sample inlets.

For capturing enzyme initial rates (Figure 1), five concentrations of lactate oxidase (1 mU/ml, 5 mU/ml, 10 mU/ml, 20 mU/ml, 50 mU/ml) were assayed with 5 mM lactate, in the buffer listed above as well as buffers containing no BSA and 0.1% of the detergents Tween20 and Triton-X-100.

For obtaining kinetic constants, lactate oxidase from *Aerococcus viridans* (product no. L9795), glucose oxidase from *Aspergillus niger* (product no. G2133) and glutamate oxidase from *Streptomyces* sp. (product no. G5921) were assayed with lactate, glucose and glutamate, respectively. Final enzyme concentrations used, in duplicates: 6 μg/ml, 1.2 μg/ml, 240 ng/ml, 48 ng/ml, 9.6 ng/ml, 0 ng/ml. Final substrate concentrations used (inside reaction chambers): 100 mM, 33.33 mM, 11.11 mM, 3.7 mM, 1.23 mM, 412 μM, 137 μM, 47 μM, 15 μM, 5 μM, 1.7 μM, 0 μM.

### Enzyme assays in microplates

For kinetic constants calculations, final concentrations of enzymes and substrates used were the same as for enzymatic assays in the Biomark HD system (see above). The assays were performed in low-volume 384-well black flat bottom plates (Greiner), reaction volume was 20 μl. The assay buffer contained 20 mM HEPES pH 7.4, 50 μM Ampliflu Red, 0.1 U/ml HRP. The assays were started by addition of substrate to the buffer-enzyme mix, the readouts were carried every 30 seconds for 30 minutes with an excitation filter of 544 nm and emission filter of 590 nm in a FLUOstar Omega microplate reader (BMG Labtech, Germany).

### Standard curves

For obtaining a resorufin standard curve for the Fluidigm system nine resorufin dilutions were prepared in 20 mM HEPES buffer, pH 7.4 with 0.11 μM fluorescein, and pipetted in respective sample inlets, together with a no resorufin control. Buffer containing 20 mM HEPES pH 7.4, 1.1 μM fluorescein, 0.11% Tween 20, 0.11% Triton X 100, 0.11 mg/ml BSA was placed in the assay inlets. Due to the all vs. all mixing of samples and assays inside the chip this results in multiple technical replicates for each concentration. Final resorufin concentrations in the microfluidic chambers were 10, 5, 2.5, 1.25, 0.625, 0.313, 0.156, 0.079, 0.04 and 0 μM.

For obtaining a resorufin standard curve using the FLUOstar Omega microplate reader nine resorufin dilutions were prepared in triplicates in 20 mM HEPES buffer, pH 7.4, together with a no resorufin control. Final resorufin concentrations were 12.5, 6.25, 3.13, 1.56, 0.78, 0.39, 0.195, 0.097, 0.049 and 0 μM. Measurements were obtained using an excitation filter of 544 nm and emission filter of 590 nm.

### Data analysis

Data obtained from the Fluidigm instrument was analyzed as follows. After a finished run, corners of the chip were adjusted manually in Fluidigm’s Real-Time PCR Analysis Software and the quantification of fluorescent intensities in the ROX and FAM channels were carried automatically by the software. The run data containing fluorescent intensity values were exported in a *.csv format. Two input files describing sample layout on both “assay” and “sample” sides were manually created. Custom scripts in R version 3.4.4 (www.r-project.org) were used to analyze data. Briefly, the resorufin fluorescence (ROX channel) was background-subtracted and converted to product concentration by using a standard curve. Regression models were fitted to the linear range of the data and the slope was calculated to obtain resorufin production per minute (μM min^−1^). The enzyme specific activity (μmol mg^−1^ min^−1^) was finally calculated using a reaction volume of 8.87 nl and the protein concentration used in each reaction. Scripts and raw data is available at GitHub (https://github.com/EngqvistLab/biomark_assays).

Data obtained from the FLUOstar Omega plate reader was analyzed in the same manner as data from the Fluidigm instrument. The only difference being that fluorescence intensities did not have to be extracted and that the reaction volumes were 20 μl.

## Acknowledgements

MKME thanks Daniel Cook, Anette Støttrup, and Daniel Foldager Larsen for initial discussions on the doability of using the Biomark system for enzyme assays. The authors thank Daniel Dancer for crucial technical assistance and Petter Woll for graciously granting access to his Biomark HD system, thus enabling these experiments to be carried out.

## Author contributions

MKME conceptualized the project. ER and MKME designed experiments. ER carried out experiments. MKME conceived and implemented the data analysis pipeline. ER and MKME performed data analysis. ER wrote the draft manuscript. ER and MKME carried out revisions on the initial draft and wrote the final version.

## Data availability

All raw data and computer code to process these are made freely available for re-use (https://github.com/EngqvistLab/biomark_assays).

## Supplementary data

**Supplementary table 1.** Fluorescent filters available in the Fluidigm system.

| Excitation filter | Emission filter |
| --- | --- |
| 475-41 nm | 525-25 nm |
| 530-16 nm | 570-30 nm |
| 575-31 nm | 630-30 nm |
| 632-27 nm* | 700-30 nm* |

\*optional filter

**Supplementary figure 1.**
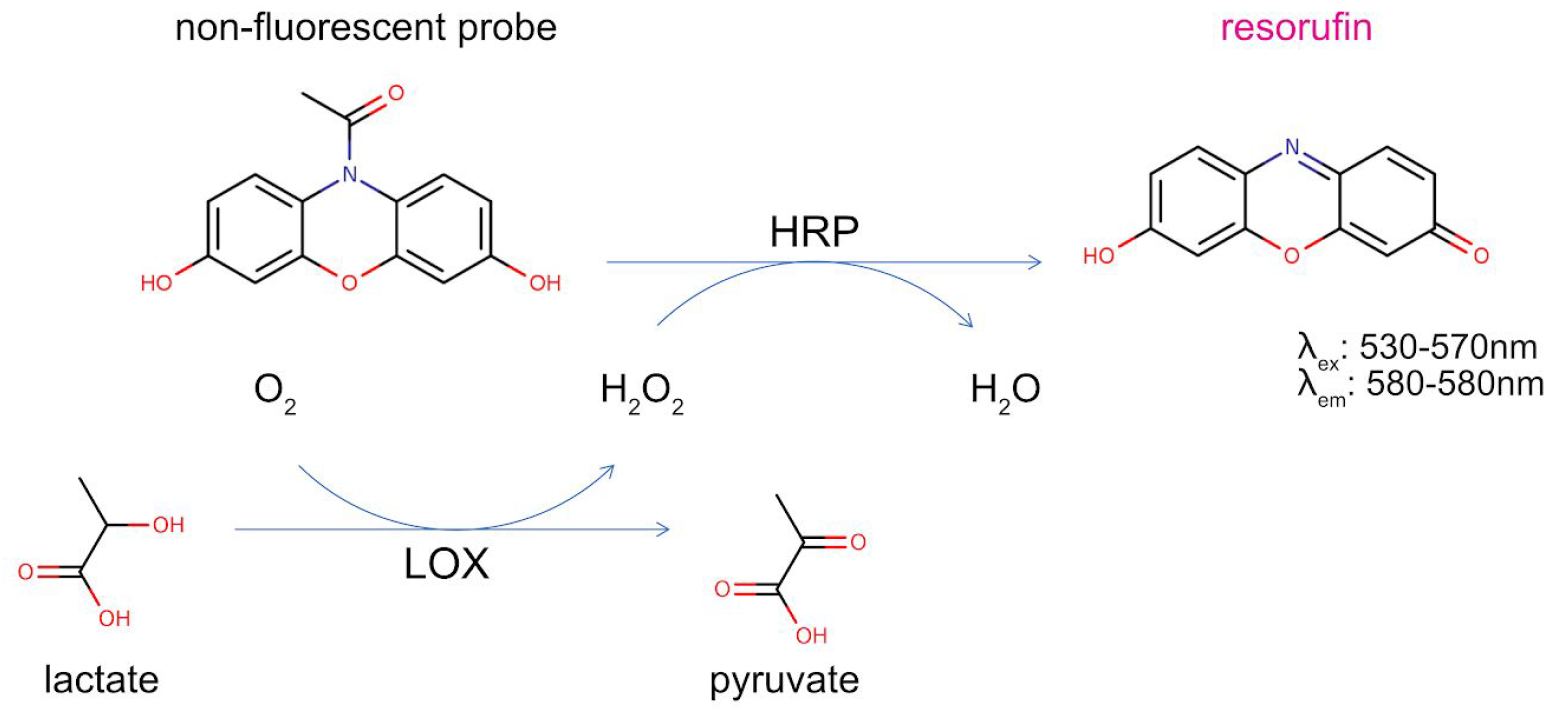
Reaction scheme of fluorescent assay for hydrogen peroxide detection on example of lactate oxidase. Lactate is oxidised to pyruvate by lactate oxidase (LOX), generating hydrogen peroxide which reacts with a non-fluorescent probe AmplifluRed in the presence of horseradish peroxidase (HRP) to produce the fluorescent oxidation product, resorufin.

**Supplementary figure 2.**
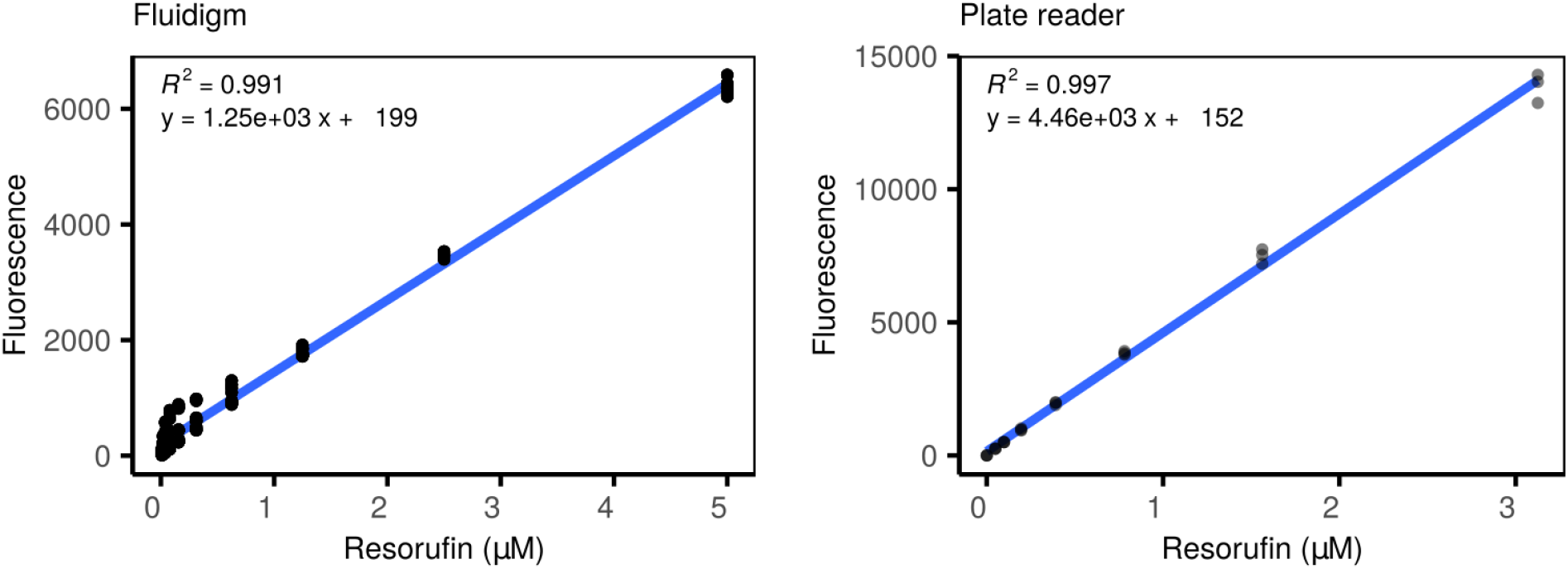
Resorufin fluorescence in the Fluidigm system and microplate reader. Represented are ranges for linear responses captured in the two systems, subsequently used as resorufin standard curves.

**Supplementary figure 3.**
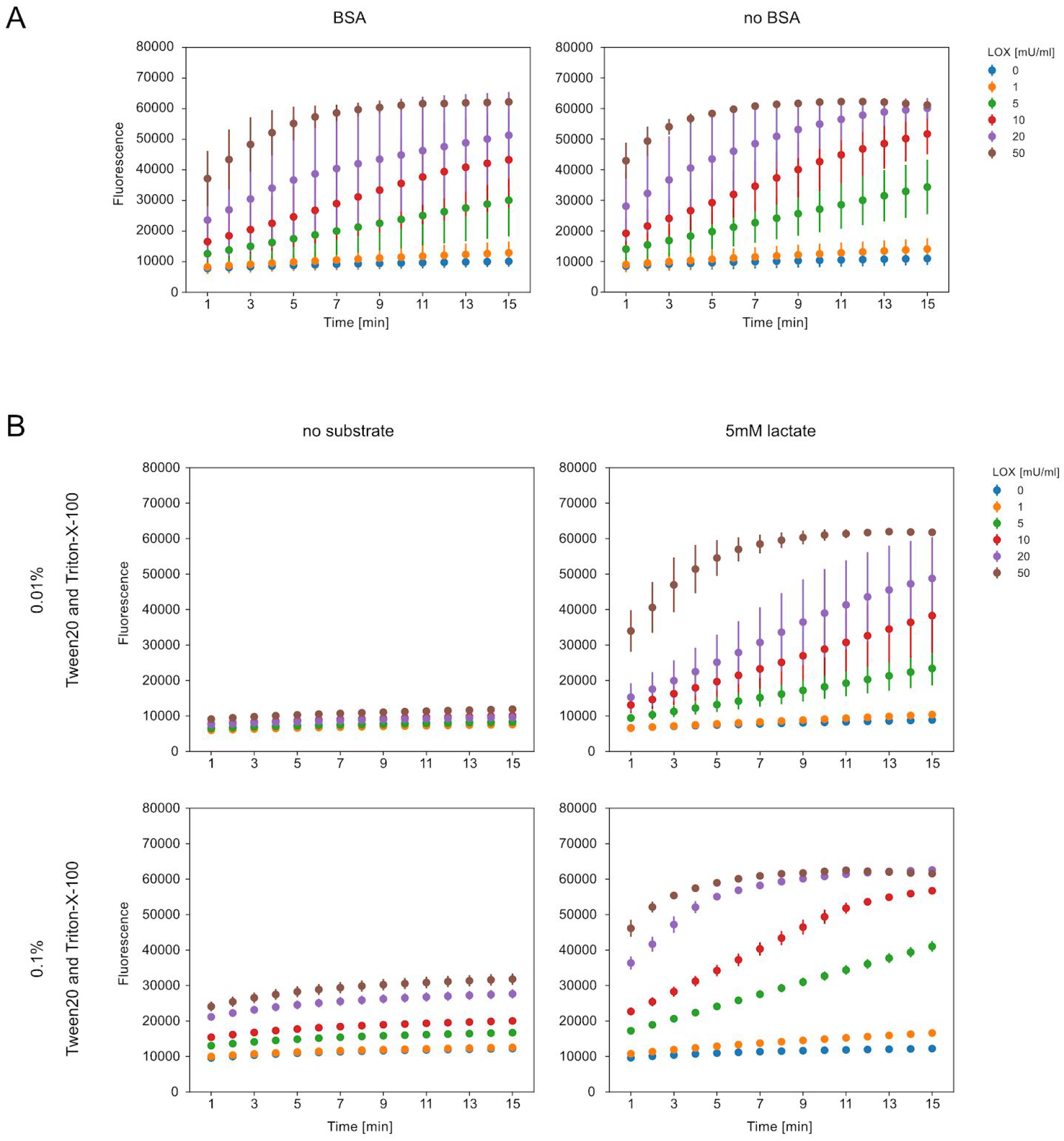
Influence of buffer composition on fluorescent readout in the Fluidigm system. (A) Fluorescent readouts for samples in buffers with and without addition of 0.01mg/ml BSA (final concentration), in presence of 5mM lactate. (B) Fluorescent readouts for samples in buffers with 0.01% and 0.1% concentration of detergents (Tween20 and Triton-X-100). Lactate oxidase (LOX) was used in these experiments. Error bars represent standard deviation of the data obtained with four replicates.

**Supplementary figure 4.**
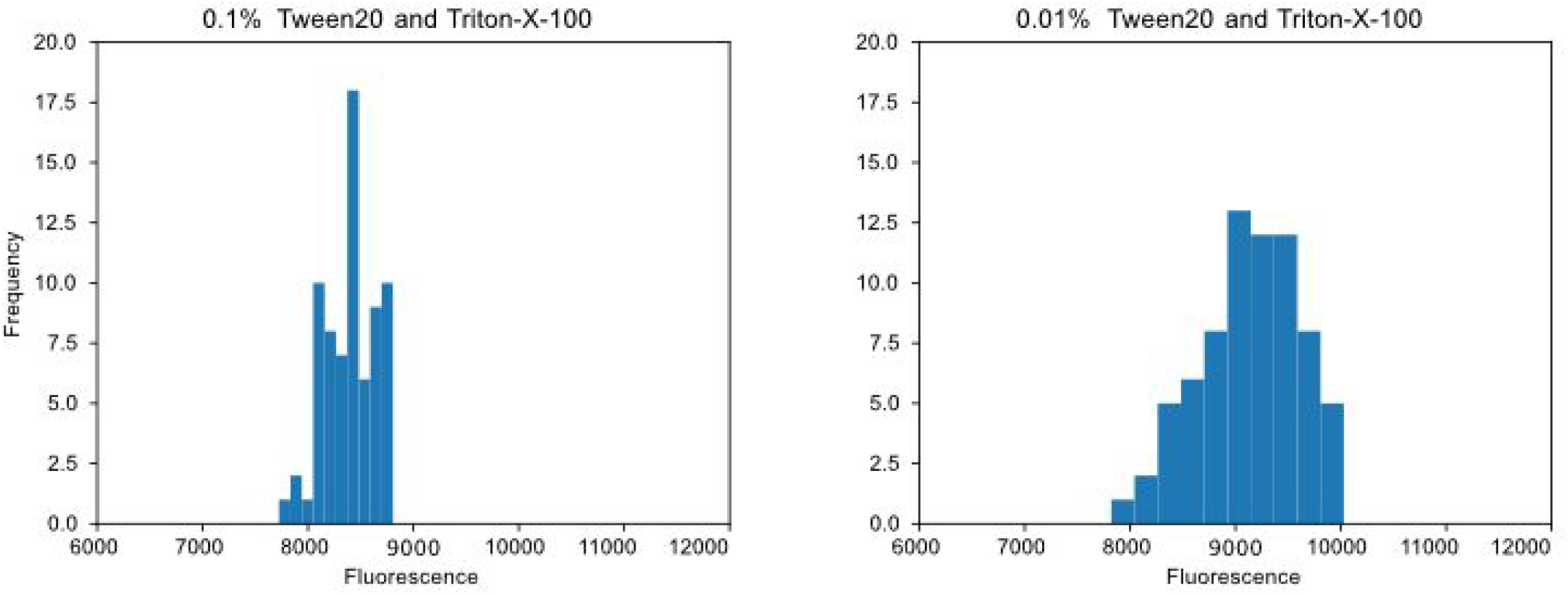
Distribution of fluorescent signal of the loading control fluorescein. Signals from samples containing detergents Triton-X-100 and Tween20 in the concentration of 0.1% (left) and 0.01% (right).

**Supplementary figure 5.**
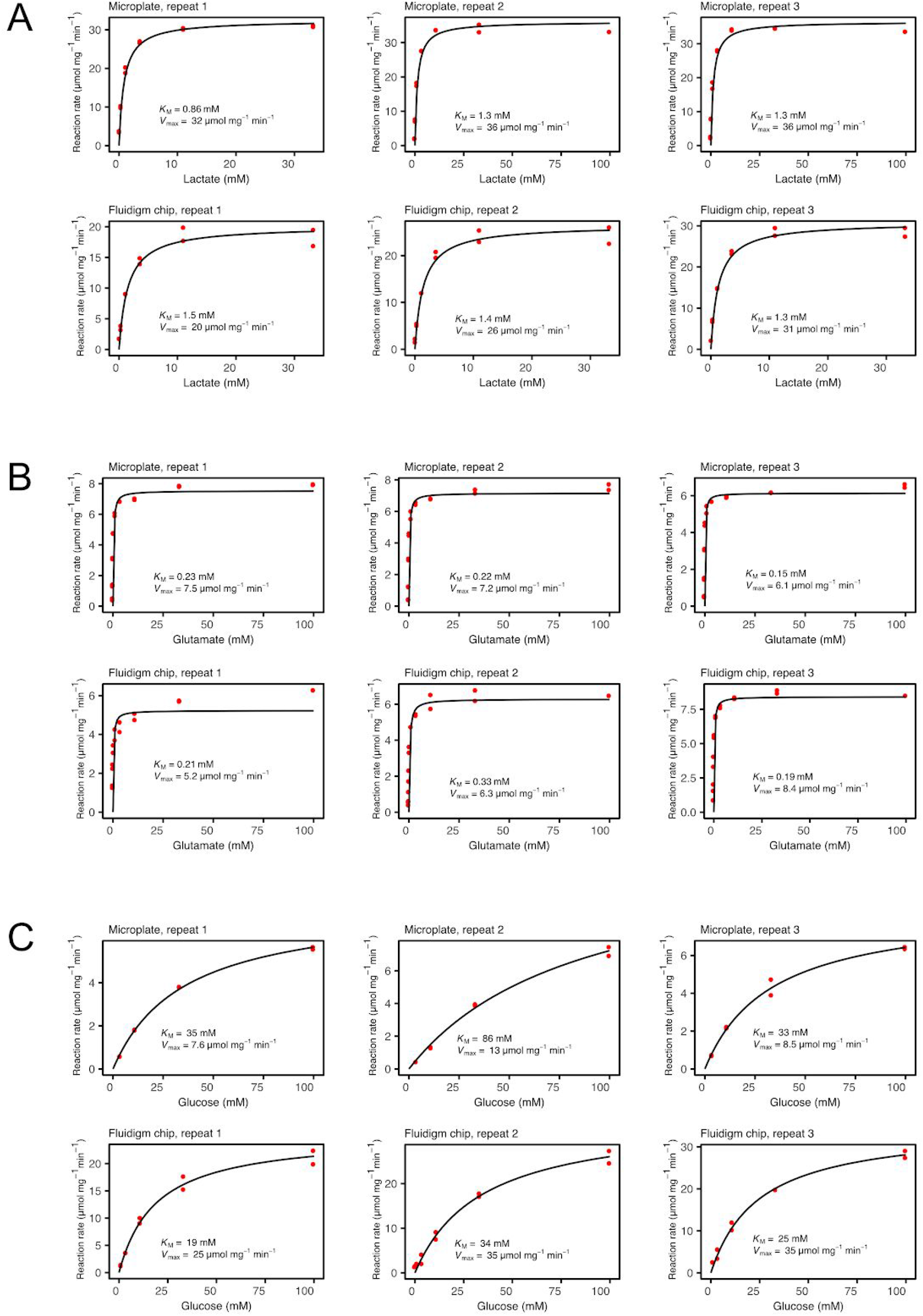
Comparison of kinetic values obtained in the Fluidigm system and a plate reader. Michaelis-Menten curves of three repeats in each system for the three tested oxidases: (A) Lactate oxidase. (B) Glutamate oxidase. (C) Glucose oxidase.

